# Single cell transcriptomics reveals distinct effector profiles of infiltrating T cells in lupus skin and kidney

**DOI:** 10.1101/2021.10.19.464575

**Authors:** Garrett S. Dunlap, Allison C. Billi, Feiyang Ma, Mitra P. Maz, Lam C. Tsoi, Rachael Wasikowski, Johann E. Gudjonsson, J. Michelle Kahlenberg, Deepak A. Rao

**Author notes:** These authors contributed equally to this work (J. Michelle Kahlenberg, Deepak A. Rao). <u>Corresponding Authors:</u> Deepak A. Rao, Hale Building for Transformative Medicine, Room 6002R, 60 Fenwood Road, Boston MA 02115, J. Michelle Kahlenberg, 5570A MSRB 2, 1150 West Medical Center Drive, Ann Arbor, MI 48109.

## Abstract

Cutaneous lupus is commonly present in patients with systemic lupus erythematosus (SLE) but can also exist as an isolated manifestation without further systemic involvement. T cells have been strongly suspected to contribute to the pathology of cutaneous lupus, yet our understanding of the T cell phenotypes and functions in the skin in lupus remains incomplete, and the extent to which lupus T cell infiltrates in skin resemble those in other tissue beds is unknown. Here, we present a detailed single-cell RNA sequencing profile of T and NK cell populations present within lesional and non-lesional skin biopsies of patients with cutaneous lupus. We identified multiple lymphocyte clusters, including both CD4 and CD8 T cells, NK cells, regulatory T cells, and a population of strongly interferon-responding cells that was present in patients with cutaneous lupus but absent in healthy donors. T cells across clusters from both lesional and non-lesional skin biopsies expressed elevated levels of interferon simulated genes (ISGs); however, compared to T cells from control skin, T cells from cutaneous lupus lesions did not show elevated expression profiles of activation, cytotoxicity, or exhaustion. Integrated analyses comparing skin T/NK cells to lupus nephritis kidney T/NK cells indicated that skin lymphocytes appeared less activated and lacked the expanded cytotoxic populations prominent in lupus nephritis. An integrated comparison of skin T cells from lupus and systemic sclerosis revealed similar activation profiles but an elevated ISG signature specific to cells from lupus skin biopsies. Overall, these data represent the first detailed transcriptomic analysis of the of T and NK cells in cutaneous lupus at the single cell level and have enabled a cross-tissue comparison that highlighted the stark differences in composition and activation of T/NK cells in distinct tissues in lupus.

## Introduction

Systemic lupus erythematosus (SLE) is a highly heterogenous disease, with the potential to manifest an array of pathologies across the body (1). Around 70% of individuals affected by SLE have cutaneous involvement, known as cutaneous lupus erythematosus, though such manifestations can also occur without further systemic symptoms (2). To date, no therapies specifically aimed at cutaneous lupus have been approved, and the current standard of care generally relies on topical corticosteroids and calcineurin inhibitors to attempt to address symptoms (3). A deeper understanding of the cells implicated in cutaneous lupus, both with and without broader systemic involvement, will be beneficial to the development more efficacious and targeted therapies.

T cells are suspected to play a major role in lupus pathology. Expanded populations of CD4 T follicular helper (Tfh) and T peripheral helper (Tph) cells promote B cell activation and autoantibody production (4–7). Work analyzing the role of regulatory T (Treg) cells in lupus has, on the other hand, been more contentious with conflicting results suggesting both increased and decreased presence in the disease (8–12). Single cell transcriptomic analyses of lupus nephritis (LN) kidney biopsies have suggested a role for cytotoxic T cell subsets in the kidneys of affected patients, with populations of NK cells, cytotoxic T cells, and Granzyme K+ CD8 T cells all being highly represented among lymphocytes in kidney biopsies (13). Further, histologic analyses of cutaneous lupus biopsies identified heterogenous staining of granzyme B+ T cells in cutaneous lesions (14). The extent to which T cell infiltrates within different target tissues in lupus, for example skin and kidney, demonstrate similar effector phenotypes, remains unclear.

Advances in single-cell RNA sequencing (scRNA-seq) technologies have allowed for the high-throughput generation and analysis of cellular states in disease and homeostasis (15, 16). To date, these tools have been applied to multiple rheumatic diseases, including lupus (17). Previous studies in lupus have produced single-cell catalogs of the cell states present in kidney biopsies of patients with LN and in PBMCs of patients with SLE (13, 18, 19). These scRNA-seq datasets have served as a foundation for a better understanding of the cell types relevant to lupus pathology in these tissues. A single-cell transcriptomic analysis of cutaneous lupus could likewise help to reveal insights into the pathogenesis of the disease and serve as a means to compare the role of a particular cell type, such as T cells, across multiple tissues of lupus pathology.

Here, we report the first detailed transcriptomic evaluation of the T and NK cells in cutaneous lupus lesions at the single-cell level. With paired lesional and non-lesional skin biopsies from patients with cutaneous lupus, as well as skin biopsies from healthy donors, we define and compare the T cell populations present across these samples. Further, we employ a previously published dataset of kidney biopsies in LN patients (13) to perform an integrated analysis of the T cell states present across both pathologic kidney and skin. Lastly, we perform a comparison of skin T cells between cutaneous lupus and systemic sclerosis (SSc) patients, providing a cross-disease examination of the role of T cells across different rheumatic skin pathologies (21). Combined, these analyses reveal both parallels and distinct differences between the T cells in skin and kidney in lupus, suggesting that the effects of therapeutic targeting of T cells may differ in different target tissues.

## Results

### Single cell transcriptomic identities of skin-localized T cells in cutaneous lupus

Skin biopsies were obtained from both lesional and non-lesional locations on 6 SLE patients and an additional patient with isolated cutaneous lupus, as well as 14 healthy controls (Table S1). Biopsies were dissociated and droplet-based scRNA-seq was used to obtain the transcriptomes of T cells. Following the application of quality control filters, we obtained a total of 3,499 T and NK cells for further analysis (Fig. 1A). Among these, we broadly identified populations of CD4+ T cells, CD8+ T cells, and NK cells (marked by *TYROBP* expression) (Fig. 1B). Further analysis identified 13 subclusters, which largely appear independent of any batch effects (Fig. 1C and S1).

**Figure 1.**
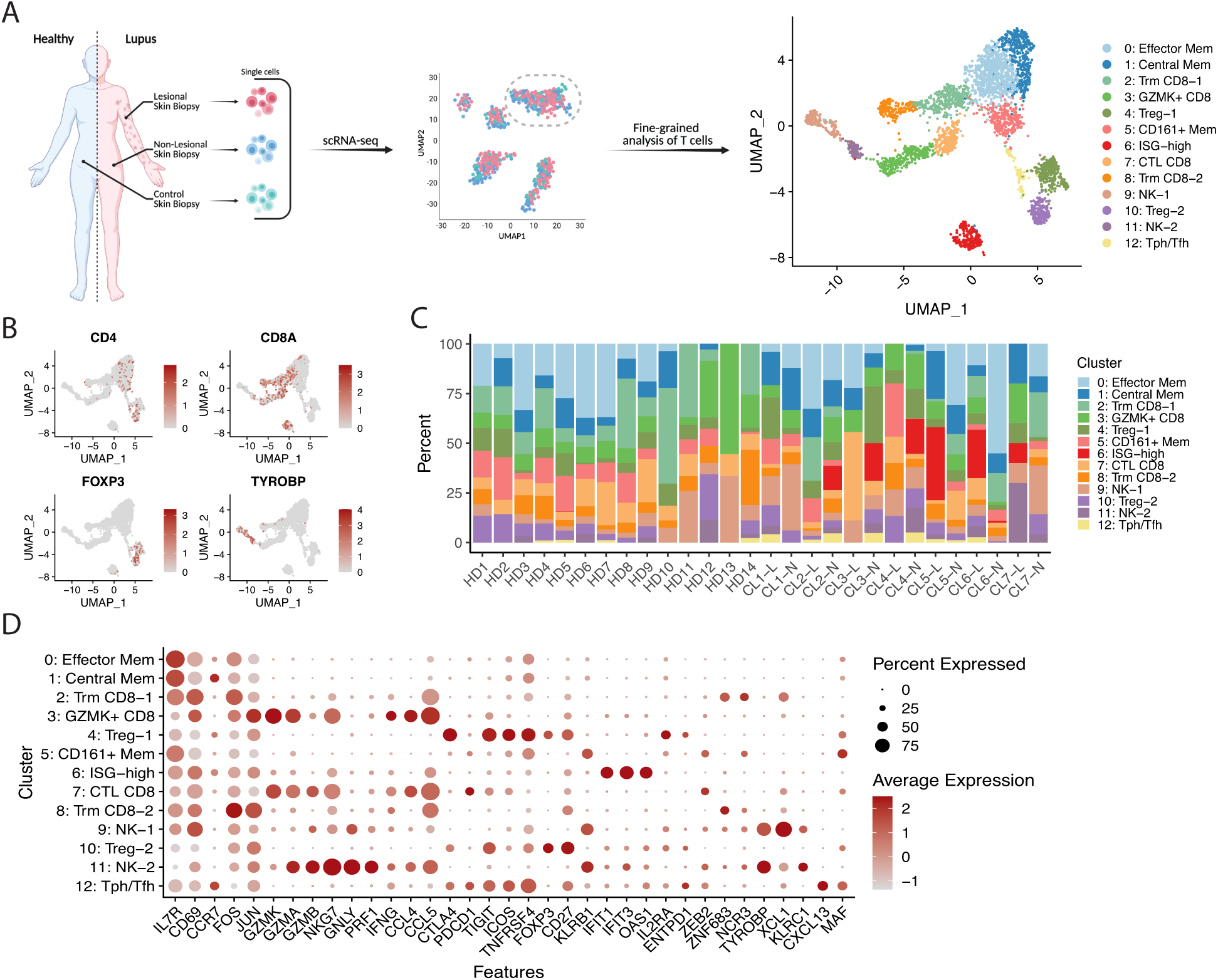
Identification of T and NK cell states in lesional and non-lesional skin biopsies of cutaneous lupus patients. **A**. Schematic diagram of study, including sample collection, initial scRNA-seq, and fine clustering of T cell subsets. **B**. UMAP plots of main T cell lineage marker expression. **C**. Bar plot of cluster assignments for captured cells in each sample. **D**. Dot plot of differentially expressed genes in each identified cluster. HD, healthy donor. CL, cutaneous lupus. L, lesional. N, non-lesional.

We next set out to define the unique transcriptional programs of each subcluster (Table S2). Across subclusters containing CD4+ cells, conventional memory T cells accounted for 3 of these clusters (#0, 1, and 5). FOXP3+ Treg cells were found across 2 clusters, both sharing expression of *CTLA4, ICOS*, and *CD27* (#4 and 10). Notably, the Tregs associated with cluster #4 were found to have stronger expression of *CTLA4, TIGIT*, and *ICOS* than those in cluster #10, while those in cluster #10 were marked by higher expression of *CD27*. The differing expression profiles of these Treg clusters suggest that those cells contained in cluster #4 belong to a more activated and suppressive subset (22). In addition, we noted a population of Tfh and/or Tph cells, defined by the absence of *FOXP3* and the expression of markers such as *CXCL13, PDCD1, ICOS*, and *MAF* (Fig. 1D).

Across the NK and CD8+ T cell clusters identified in these biopsies, tissue-resident memory CD8+ T (Trm) cells, identified by expression of markers such as *ZNF683* and *XCL1*, formed 2 clusters (#2 and 8). A population of CD8+ T cells marked by the strong expression of *GZMK* but relative absence of *GZMB* was identified (#3), consistent with a phenotype previously recognized in LN kidneys and rheumatoid arthritis synovium (13, 23). In contrast, cluster #7 expressed higher levels of *GZMB* compared to cluster #3, suggesting a cytotoxic T lymphocyte (CTL) phenotype. Clusters #3 and #7, though, were both found to similarly express the cytokine *IFNG* and chemokines *CCL4* and *CCL5*. Aside from these CD8+ clusters, 2 subclusters of NK cells were identified in the dataset (#9 and 11). While both share a similar core transcriptional program composed of markers such as *KLRB1, TYROBP*, and *NKG7*, the NK cells of cluster #9 are largely differentiated by stronger expression of *XCL1*, suggestive of a CD56^bright^ NK cell population, while those of cluster #11 overexpress *PRF1*, multiple granzyme genes, and *CCL5*, likely representing a CD56^dim^ population (24).

In addition, a cluster marked by the expression of interferon-stimulated genes (ISGs) such as *IFIT1, IFIT3, IFI44, OAS1*, and *LY6E*, among others, was identified. This population of cells appears to be a mixture of CD4+, CD8+, and CD4-CD8-cells, likely comprising cells from the other major subclusters (Fig. 1D). This ISG-enriched cluster is consistent with previously published scRNA-seq studies of the T cells in nephritic kidney, blood, and skin of SLE patients, further highlighting a conserved interferon signature across cell types and tissues in SLE (13, 19, 20).

### Similarly elevated interferon-response signature in T cells at both lesional and non-lesional sites in cutaneous lupus

We next sought to determine if certain cell subsets were differentially represented among the lesional, non-lesional, and control samples obtained by this study. Of the 3,499 T and NK cells sequenced, 2,116 and 1,383 were collected from cutaneous lupus patients and healthy controls, respectively. In total, 687 cells were obtained from the lesional biopsies of these lupus patients, and 1,429 were sequenced from their paired non-lesional samples. Accounting for differences in cell numbers for each sample type, we identified the ISG-high cluster to have greater representation in the lupus patient samples compared to healthy controls (Fig. 2A and B).

**Figure 2.**
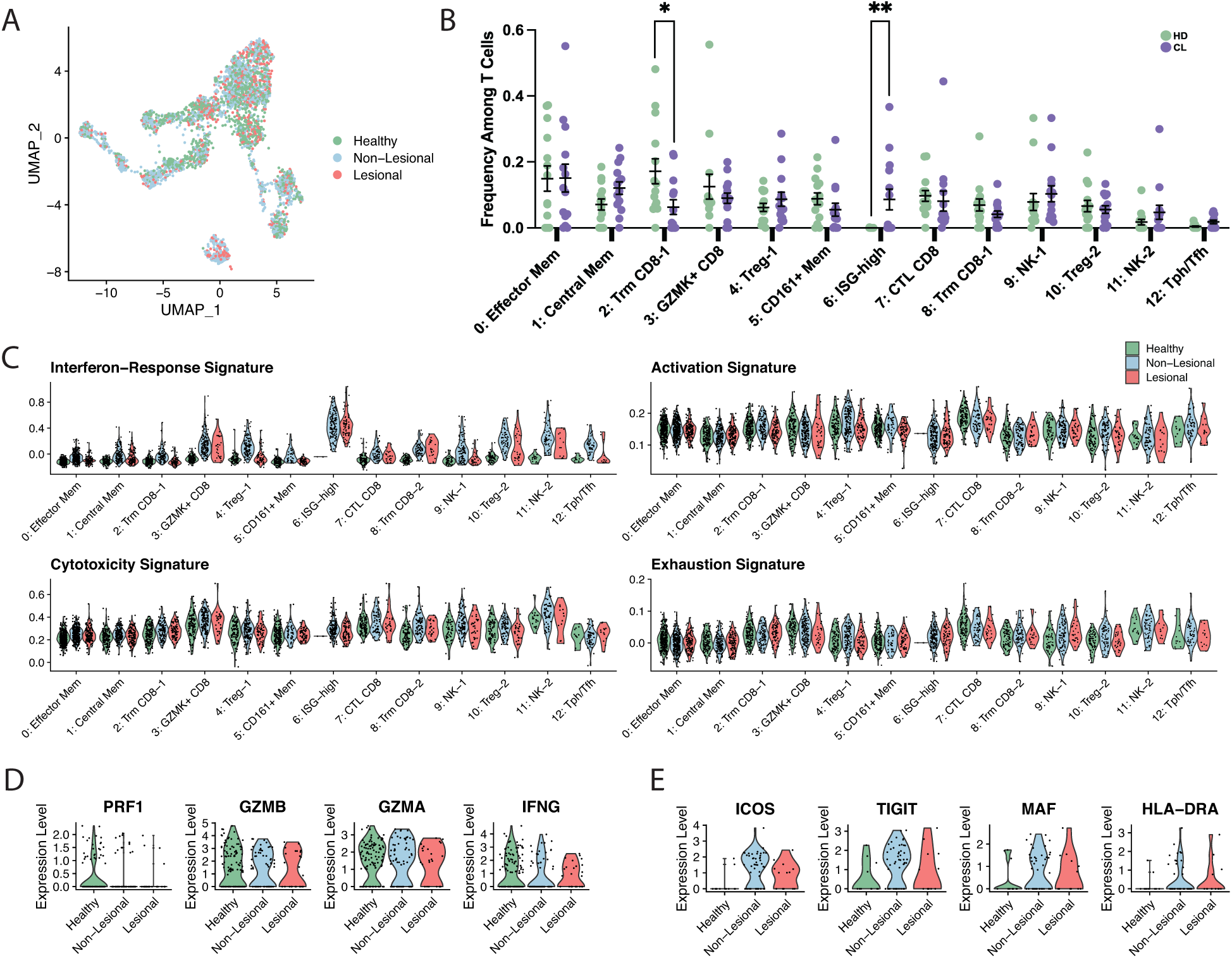
Examination of differences between lesional, non-lesional, and healthy skin biopsies. **A**. UMAP plot of cells colored by sample type. **B**. Comparison of the frequencies of T cells per cluster between cutaneous lupus (CL) patients and healthy donors (HD). **C**. Violin plots of signature scores between healthy, lesional, and non-lesional cell components of each cluster. **D**. Violin plots of the expression of select markers between healthy, lesional, and non-lesional cell components of the CTL CD8 cluster. **E**. Violin plots of the expression of select markers between healthy, lesional, and non-lesional cell components of the Tph/Tfh cluster. Mean ± SEM shown. ^*^p<0.05, ^**^p<0.01 by Mann-Whitney U test in (B).

The cluster of ISG-high T cells was nearly exclusively represented by cells from the cutaneous lupus samples, including cells from both lesional and non-lesional samples (Fig. S2). In addition, an elevated ISG transcriptional signature was seen across clusters in both the lesional and non-lesional samples compared to controls (Fig. 2C). This is consistent with previous studies that have identified a conserved elevation in type I interferon-regulated genes across patients with cutaneous lupus erythematosus and SLE (25–27).

We next sought to use the transcriptomic data to compare the functional status of T cells in cutaneous lupus and healthy skin biopsies. For this effort, we generated activation-, cytotoxicity-, and exhaustion-relevant gene lists and calculated signature scores for lesional, non-lesional, and healthy controls (Table S3). Surprisingly, T cells from cutaneous lupus lesions and non-lesional sites appeared quite similar to T cells from control skin across these measures, suggesting a lack of wide-scale activation in T cells within cutaneous lupus skin lesions (Fig. 2C).

A focused analysis of the expression of key cytotoxic genes specifically within the CTL cluster similarly revealed no expression differences (Fig. 2D). In contrast, a focused analysis on the Tph/Tfh cell cluster, a population of cells strongly implicated in pathologic T/B cell interactions in SLE, revealed some differences. Notably, we found an upregulation of costimulatory genes such as *ICOS* and *TIGIT* in Tph/Tfh cells from cutaneous lupus patients compared to their healthy control counterparts, and further noted an upregulation of HLA-DRA and the transcription factor *MAF*, which has been demonstrated to promote Tph/Tfh cell function (Fig. 2E) (13).

Altogether, these data highlight the systemic nature of detection of interferon response genes in T cells of the skin and suggest a potential increase in activity of the B cell-helping Tph/Tfh cells, but otherwise indicate limited features of activation or cytotoxicity in skin-localized T cells in cutaneous lupus.

### Low cytotoxicity and effector signatures in T cells from cutaneous lupus compared to lupus nephritis

To better understand our results in the broader context of lupus, we sought to define conservation and potential differences across T cells found at different tissue sites in SLE. To accomplish this, we performed an integration of our dataset with T cells from a previously published scRNA-seq dataset of immune cells obtained from kidney biopsies of patients with LN (25). Within this dataset, we isolated 1719 T and NK cells from 24 patients with LN. Following integration of these datasets using canonical correlation analysis (CCA), we produced a single unified visualization of data from both tissues (Fig. 3A). Cells of the same subset largely clustered together irrespective of tissue origin, and all of the T cell types were found in both tissues, including Trm CD8+, Tregs, cytotoxic CD8+, Tph/Tfh cells, and NK cells (Fig. 3B,C).

**Figure 3.**
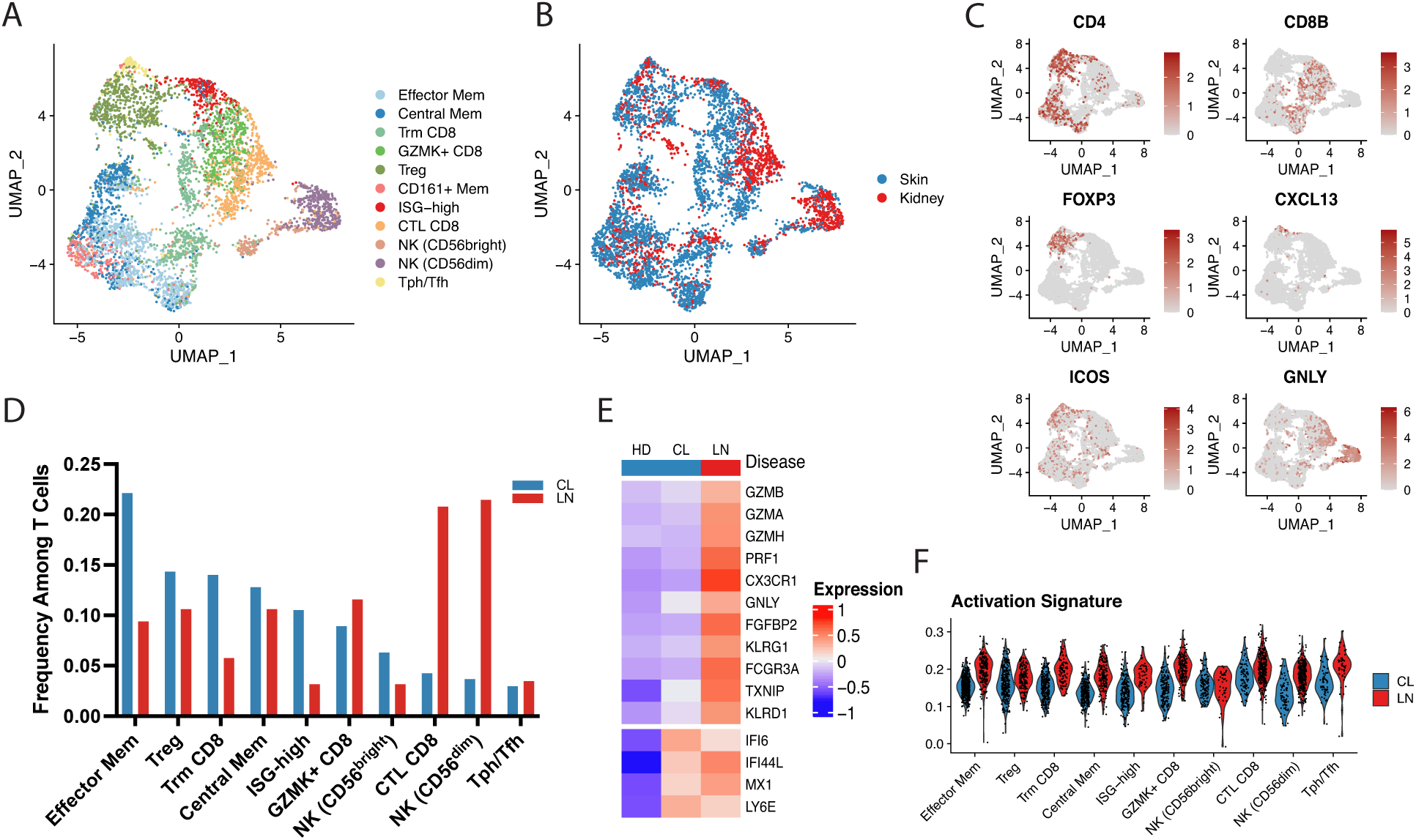
Integration of cutaneous lupus and lupus nephritis single-cell datasets reveals decreased cytotoxic profiles in the skin T cells. **A**. UMAP plot of the cluster identifications resulting from canonical correlation analysis (CCA) integration. **B**. UMAP plot of the location of cells from each study. **C**. Representative UMAP plots of select major lineage markers. **D**. Comparison of the frequencies of T cells per cluster between cells from cutaneous lupus (CL) and lupus nephritis (LN) datasets. **E**. Scaled heatmap of cytotoxic genes (top) and select interferon-stimulated genes (bottom). **F**. Violin plot of the activation signature scores between CL and LN cells for each cluster.

When comparing representation of these cell states across tissues, we noted an increased proportion of memory T cells in the skin of cutaneous lupus patients, including CD4 memory and CD8 Trm subsets, along with a larger percentage of cells in the ISG elevated cluster, when compared to kidney samples of LN patients. Conversely, we observed a strongly increased representation of cells with cytotoxic function, including CD8+ and NK clusters, in samples obtained from the kidneys of LN patients (Fig. 3D).

Gene-level examination of cytotoxic marker expression between cutaneous lupus and LN further highlighted this difference between T cells from the different tissues, with LN T cells having increased expression of genes associated with cytotoxicity, including *GZMB, GZMH, PRF1*, and *GNLY* (Fig. 3E). In comparison, ISGs showed similar expression levels between cutaneous lupus and LN T cells, indicative of the systemic interferon response in lupus. Along with an increase in cytotoxicity genes, T cells from LN kidneys also showed an increased activation signature score across clusters compared to T cells from skin (Fig. 3F).

### Elevated IFN signatures in skin-localized T cells from lupus compared to systemic sclerosis

To further extend our exploration of the T cells present in cutaneous lupus, we sought to compare our data to T cells from the skin of another rheumatic disease. A recent study focused on systemic sclerosis (SSc) obtained skin biopsies from 27 patients with SSc and an additional 10 healthy controls (21, 28). After isolating T cell transcripts from this data, we integrated the dataset with our dataset of cutaneous lupus T cells, providing a unified visualization of both sets in the same UMAP space (Fig. 4A). Similar to our results upon integrating data from skin T cells with data from kidney T cells in lupus, we observed the presence and co-clustering of all major cell types described above in T cells from both systemic sclerosis skin and lupus skin samples (Fig. 4B).

**Figure 4.**
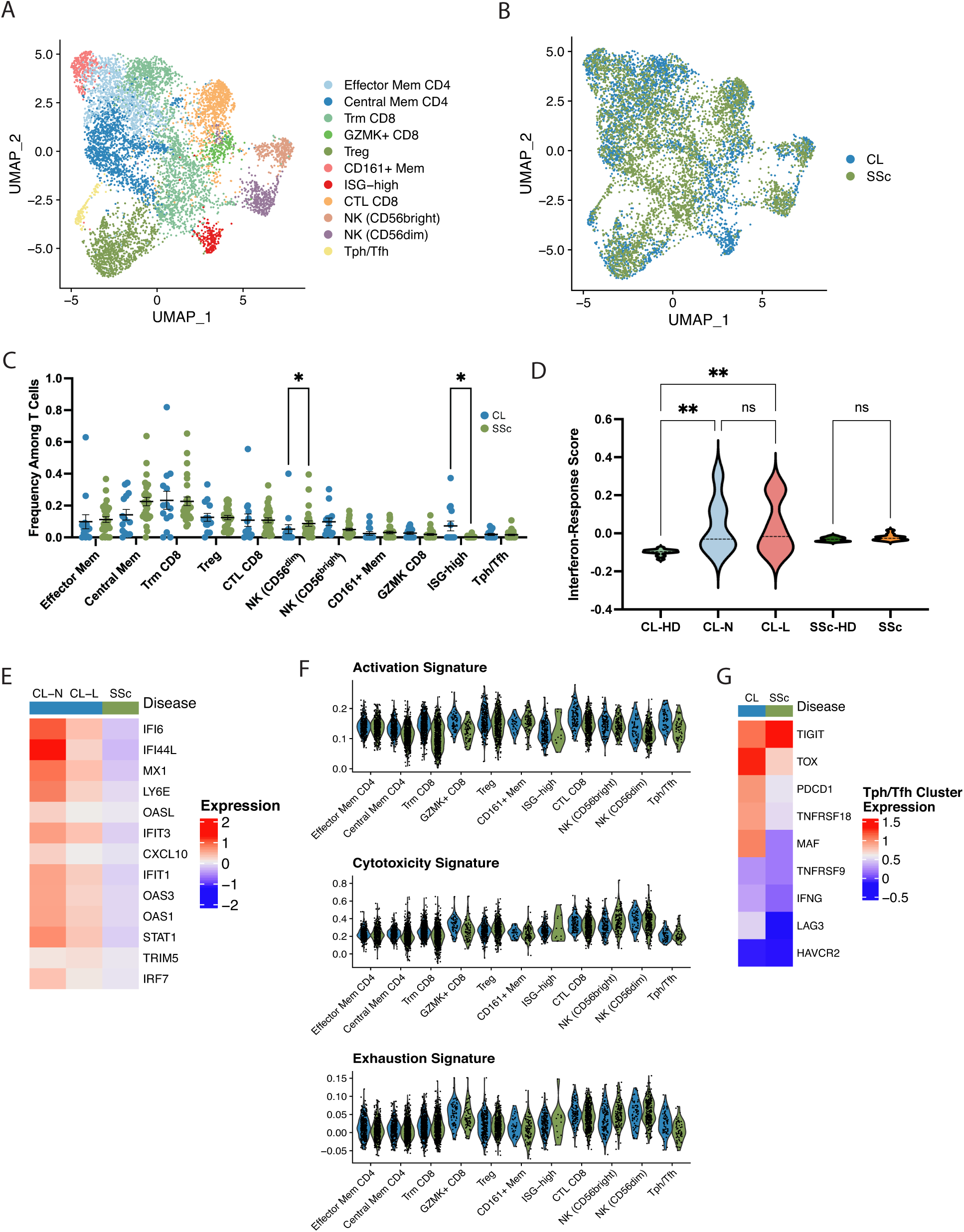
Integration of cutaneous lupus and systemic sclerosis single-cell datasets demonstrates selective ISG upregulation in lupus. **A**. UMAP plot of the cluster identifications resulting from CCA integration. **B**. UMAP plot of the location of cells from each study. **C**. Comparison of the frequencies of T cells per cluster between cells from cutaneous lupus (CL) and systemic sclerosis (SSc) datasets. **D**. Violin plot of interferon response signatures between study and sample types. **E**. Scaled heatmap of interferon-stimulated genes. **F**. Violin plots of signature scores between CL and SSc cells for each cluster. **G**. Scaled heatmap of select activation and exhaustion markers in the Tph/Tfh cluster. Mean ± SEM shown. ^*^p<0.05, ^**^p<0.01 by Kruskal-Wallis test in (C) and (D).

Comparing the distribution of cells from each disease for each cluster, we noted a deficiency of cells from SSc samples in the cluster associated with the strongest ISG signature (Fig. 4C). Examination of the sample groups and healthy controls revealed that while there is a significant increase in ISGs comparing T cells from control skin and T cells from cutaneous lupus lesional and non-lesional sites, no such difference exists between SSc samples and their respective healthy controls (Fig. 4D). A heatmap analysis of the expression patterns of multiple ISGs in cutaneous lupus and SSc T cells further corroborated this finding, suggesting that IFNs more strongly influence the cutaneous T cell response in lupus than in SSc (Fig. 4E).

We them aimed to profile differences in the activation and effector function of T cells between cutaneous lupus and SSc. Through an examination of signature scores, our analysis noted no differences in these scores between the cutaneous lupus and SSc components of each identified cluster (Fig. 4F). Lastly, we sought to compare the Tph/Tfh cluster, which is marked by *CXCL13*+ CD4 T cells in cutaneous lupus and SSc. While ISGs represented many of the most upregulated genes in cutaneous lupus, we also noticed an upregulation of multiple activation and exhaustion genes. Notably, we found higher levels of *PDCD1, TOX, LAG3, TNFRSF18*, and *MAF* on average in T cells from cutaneous lupus samples compared to SSc samples (Fig. 4G).

Together, these results emphasize the strength of interferon-response in lupus, even in comparison with another rheumatic disease, and suggest an increased activity of B cell-helping T cell subsets in cutaneous lupus compared to SSc.

## Discussion

This study represents the first in-depth examination of skin-localized T cells in cutaneous lupus using single-cell transcriptomics. Our analysis revealed deep transcriptional heterogeneity within the T cells collected from the skin of these patients, including subsets of CD4 and CD8 memory T cells, Tregs, cytotoxic T cells, helper T cells, and others. One of the defining characteristics in our comparison of T cells from lupus patients and healthy donors is the significant upregulation of ISGs. Though the role of IFN is increasingly well-understood in the pathogenesis of lupus (28), this study furthers the notion of systemic effects of this signaling that can be detected across tissues. Our results demonstrate robust upregulation of ISGs in T cells of both the kidney and skin of lupus patients, far exceeding that in T cells from both healthy skin and skin from SSc patients.

Aside from strong differences related to interferon response, we noted surprisingly little transcriptional evidence of increased activation or effector function engagement when comparing T cells from cutaneous lupus skin to those from healthy skin. In contrast, when comparing T cells from the kidneys of lupus patients with T cells from the skin of lupus patients, our analysis suggests that the T cell infiltrates differ substantially at these two tissue sites. We noted a marked increase in T cells associated with cytotoxic function, and likewise observed an increase in expression of effector function-related genes, in T cells from LN kidneys. These findings suggest a different role of T cells in the pathology of lupus in disparate tissues, whereby T cells in organs such as the kidney may mediate cytotoxic effects that contribute to tissue injury, while T cells in the skin may contribute alternative functions, including B cell help, or may be primarily bystanders.

There are several limitations that raise caution about this interpretation of the data. First, the total number of lesional T cells analyzed is limited. Second, it is possible that T cell activation signatures are downregulated during the longer processing time required for isolation of T cells from skin. However, the robust detection of ISGs suggests that at least some of the disease-associated transcriptomic signatures are retained during processing. Third, it is possible that activated T cells are inefficiently collected or preferentially depleted during tissue processing, or that the activated cells represent a small minority of the total cells that is not well visualized in our analyses. Further analyses by complementary approaches will be helpful to address these considerations. Yet taken together, the striking differences between transcriptomes of T cells from kidney and skin appear to suggest substantial differences in their functions.

Notably, our analysis of the Tph/Tfh cells of cutaneous lupus patients suggests an increased activation of Tph/Tfh cells obtained from lesional and non-lesional lupus skin compared to control skin or SSc skin. These cells may have a role in directing B cell responses within the skin. However, the frequency of Tph/Tfh cells in cutaneous lupus lesions is low and not clearly different from controls; thus, it is also possible that these cells in skin are a reflection of the activated circulating Tph/Tfh cell populations in SLE patients (29). The findings related to this cell subset and their potential role in the skin of lupus patients warrant further examination with larger populations of isolated cells.

The search for effective and targeted therapies for lupus remains difficult, and certain gaps in our knowledge of the diseases pathogenesis across tissues remain. Here, we delineate the phenotypes of T cells in cutaneous lupus, both in relation to healthy donor T cells as well as T cells from other affected organs. These results suggest a limited activation of T cells in the skin of lupus patients, especially in comparison to those of the kidney, and suggest that the pathologic roles of T cells in lupus differ depending on the target tissue. These findings may help to explain the differences in the efficacy of antimetabolite or T cell-directed therapies in lupus manifestations in skin and kidney and may influence the search for the most efficacious pathways to target to treat cutaneous lupus.

## Methods

### Subjects and Sample Collection

Skin punch biopsies were obtained from lesional (primarily back, chest, arms) and non-lesional (sun-protected hip) locations from 7 patients with active cutaneous lupus (Supplementary Table 1). A diagnosis of SLE was confirmed for 6 of 7 patients via the 2019 EULAR/ACR criteria, and the remaining patient was noted to have cutaneous involvement only. In addition, 14 healthy donors were recruited to obtain control skin punch biopsies from sun-protected skin. Sample collection for this study was approved by the University of Michigan Institutional Review Board (IRB).

### Single-cell cDNA and Library Preparation

The collected biopsies were incubated overnight in 0.4% dispase (Life Technologies) in Hank’s Balanced Saline Solution (Gibco) at 4°C as previously detailed (29). Single-cell libraries for all samples were generated using the 10x Genomics Chromium Single Cell 3’ (v3 Chemistry) protocol through the University of Michigan Genomics Core. Finally, libraries were sequenced on an Illumina NovaSeq 6000 sequencer to generate 150 bp paired end reads.

### Data Analysis

#### Processing FASTQ reads

Initial processing of the sequencing output, including quality control, read alignment to a reference genome (GRCh38), and gene quantification was performed using the 10x Genomics Cell Ranger pipeline (v3.1.0). Following the generation of barcode and UMI counts, samples were merged into a single expression matrix using the cellranger aggr pipeline.

#### Data subsetting and quality control filtering

T and NK cell clusters were subset from the above dataset of skin biopsies from cutaneous lupus patients and healthy donors in Seurat (v4.0.3). The remaining cells were then filtered to only include cells with between 200 and 3500 detected features, and those with less than 15% of reads associated with mitochondrial genes.

#### Dimensionality reduction and clustering

Cells that passed all subset and filter steps then underwent another round of principal component analysis, where the first 20 PCAs were subsequently used for downstream analysis. To ensure integration of our samples, we corrected batch effects at the level of the sample using Harmony (30). We employed Harmony using our PCA reduction with theta = 2, max.iter.cluster = 20, and max.iter.harmony = 10. After integration, cells were clustered using Louvain clustering in Seurat with a resolution of 0.6, determined through an iterative analysis of clustering results. Resulting clusters were then visualized using Uniform Manifold Approximation and Projection (UMAP) plots. Differential gene expression between clusters was determined using a Wilcoxian Rank Sum Test, and the resultant list was filtered to only include genes with a log fold-change greater than 0.25 and those which were detected in at least 50% of the population being tested.

#### Dataset integration

Inter-dataset integration was performed using canonical correlation analysis (CCA) in Seurat (31). First, the lupus nephritis and systemic sclerosis datasets were downloaded and filtered on the above quality control metrics, before being clustered. The T and NK cells were then subset out from the respective datasets. Following this, CCA integration was performed by first finding anchors between the dataset pairs, using the first 20 dimensions. Following, the *IntegrateData* function in Seurat was used to display the datasets on the same UMAP projection.

## Data Availability

The cutaneous lupus single-cell transcriptomics data analyzed in this publication is available in the Gene Expression Omnibus (GEO) database under the accession number XXXXXXXXX. The lupus nephritis analyzed during this study can be accessed within the Single Cell Portal hosted by the Broad Institute using the study ID SCP279.

The systemic sclerosis data analyzed during this study is available in the Gene Expression Omnibus database under the accession numbers GSE138669 and GSE181957.

## Author Contributions

Conceived and designed the analysis: GSD, JMK, DAR.

Collected the data: ACB, MPM, RW, LCT, JEG.

Performed the analysis: GSD, FM.

Wrote the manuscript: GSD, DAR.

Supervised the work: JMK, DAR.

## Acknowledgements

We would like to thank the lupus and systemic sclerosis patients that contributed samples to this study. This work has been supported in part by funding from the Burroughs Wellcome Fund Career Award in Medical Sciences (DAR); the Taubman Institute via Innovative Program (JMK, JEG); Parfet Emerging Scholar (JMK) funds; the National Institute of Health (NIH): K08-AR078251 (ACB), K01-AR072129 (LCT), P30-AR075043 (JEG), R01-AI130025 (JEG), R01-AR071384 (JMK), K24-AR076975 (JMK), K08-AR072791 (DAR), and P30-AR070253 (DAR); and the Lupus Research Alliance (JMK). Additionally, ACB, LCT and JEG are supported by the Dermatology Foundation, and LCT is supported by the Arthritis National Research Foundation and National Psoriasis Foundation.

---

**Supplementary Figure 1.**
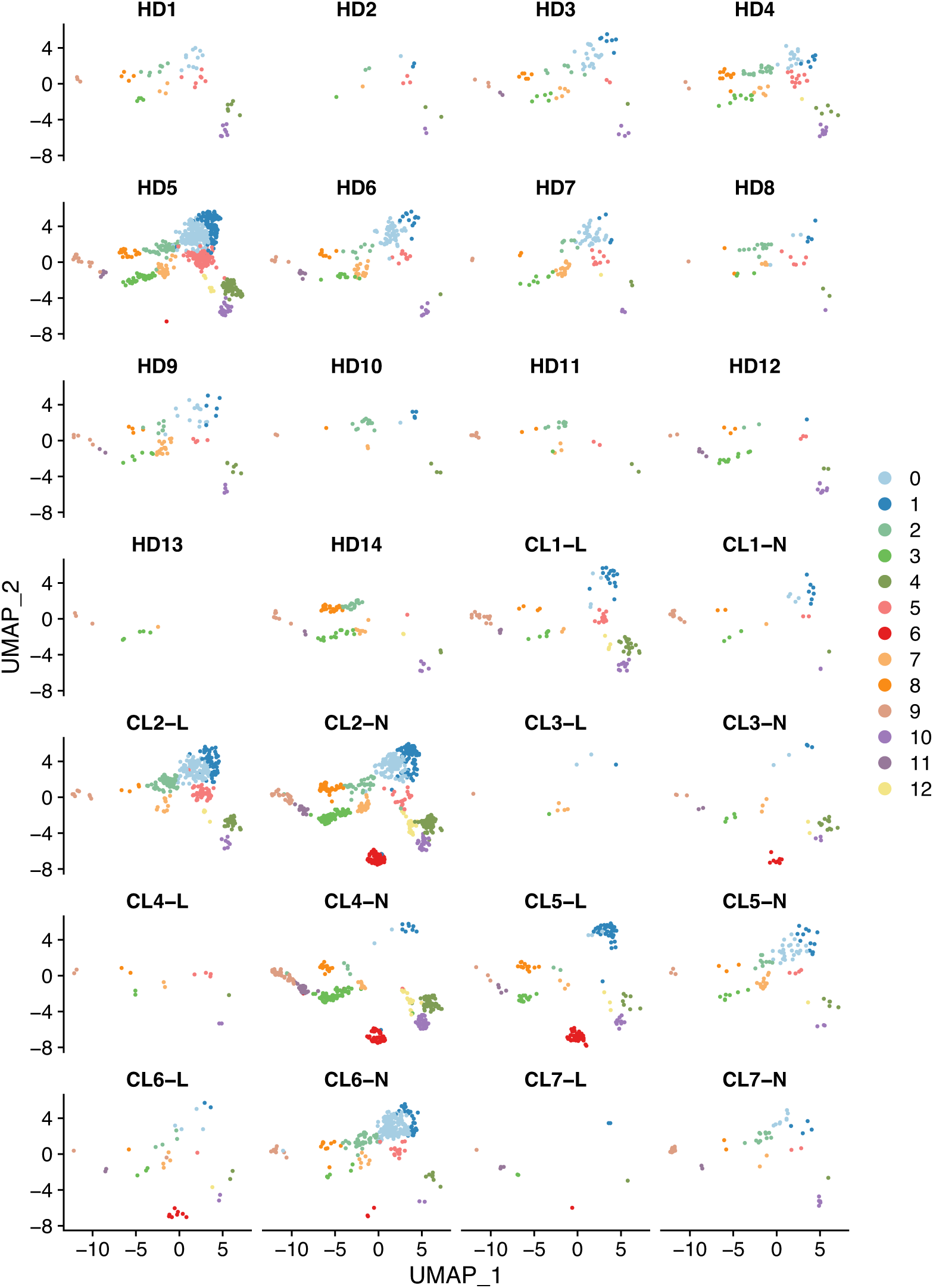
UMAP plot locations of the cells from each sample.

**Supplementary Figure 2.**
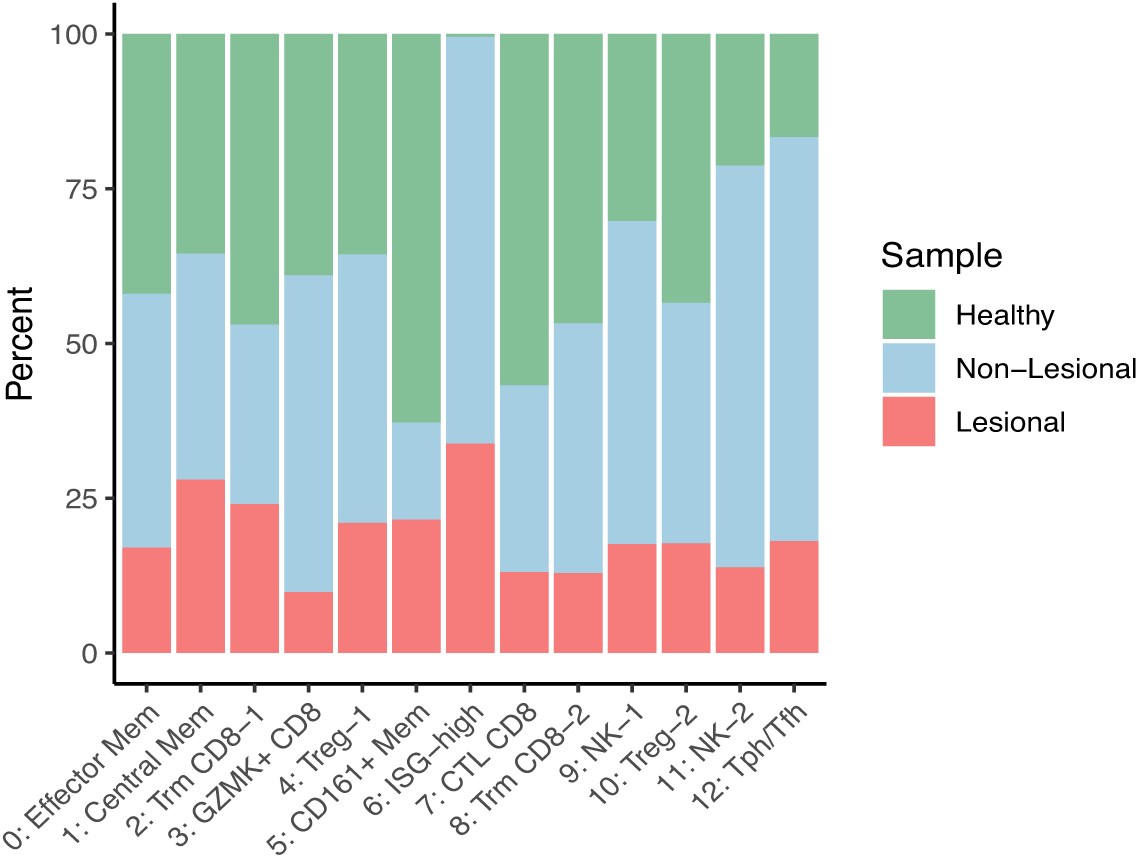
Barplot of the healthy, non-lesional, and lesional components of each cluster.

**Supplementary Figure 3.**
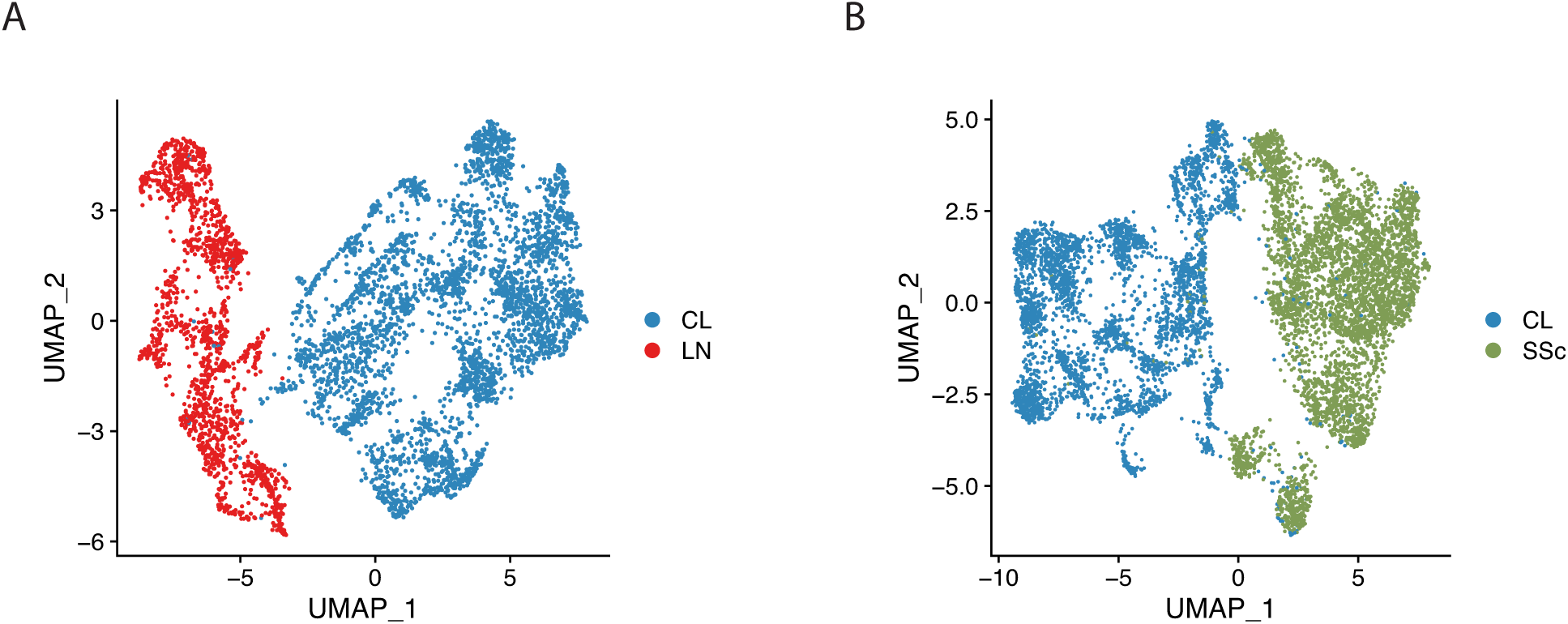
**A**. UMAP plot of the merged cutaneous lupus (CL) and lupus nephritis (LN) datasets before CCA integration. **B**. UMAP plot of the merged CL and systemic sclerosis (SSc) datasets before CCA integration.

[See uploaded file]

**Supplementary Table 1**. Clinical characteristics of cutaneous lupus patients included in this study.

[See uploaded file]

**Supplementary Table 2**. Genes differentially expressed across T/NK clusters in skin biopsy samples.

[See uploaded file]

**Supplementary Table 3**. Signature gene lists for activation, cytotoxicity, exhaustion, and interferon response.

